# *De novo* amyloid peptides with subtle sequence variations differ in their self-assembly and nanomechanical properties

**DOI:** 10.1101/2023.05.08.539858

**Authors:** Hannah G. Abernathy, Jhinuk Saha, Lisa K. Kemp, Parvesh Wadhwani, Tristan D. Clemons, Sarah E. Morgan, Vijayaraghavan Rangachari

## Abstract

Proteinaceous amyloids are well known for their widespread pathological roles but lately have emerged also as key components in several biological functions. The remarkable ability of amyloid fibers to form tightly packed conformations in a cross β-sheet arrangement manifests in their robust enzymatic and structural stabilities. These characteristics of amyloids make them attractive for designing proteinaceous biomaterials for various biomedical and pharmaceutical applications. In order to design customizable and tunable amyloid nanomaterials, it is imperative to understand the sensitivity of the peptide sequence for subtle changes based on amino acid position and chemistry. Here we report our results from four rationally-designed amyloidogenic decapeptides that subtly differ in hydrophobicity and polarity at positions 5 and 6. We show that making the two positions hydrophobic renders the peptide with enhanced aggregation and material properties while the introduction of polar residues in position 5 dramatically changes the structure and nanomechanical properties of the fibrils formed. A charged residue at position 6, however, completely abrogates amyloid formation. In sum, we show that subtle changes in the sequence do not make the peptide innocuous but rather sensitive to aggregation, reflected in the biophysical and nanomechanical properties of the fibrils. We conclude that tolerance of peptide amyloid for subtle changes in the sequence should not be neglected for the effective design of customizable amyloid nanomaterials.

## 1 Introduction

Amyloids are highly ordered self-assembling proteins characterized by their long, fibrous cross β-sheet structure^1^ and are known to have both pathological and functional implications in various organisms. While amyloid structures of some proteins have been established as key species involved in neurodegenerative diseases such as Alzheimer’s and Parkinson’s diseases, Amyotrophic lateral sclerosis (ALS), and Type-II diabetes, ^2-5^ they have also been found to play functional roles in bacterial biofilm formation,^6^ and melanin formation in mammals.^7^ Biofilms, perhaps the best example of functional amyloids, are proteinaceous extracellular fibers made up of Curli, which is orchestrated by a family of six proteins, Csg ABDEFG transcribed by the *Csg* operon^6^, with CsgA and CsgB specifically responsible for Curli formation.

The underlying commonality among amyloids is a cross β-sheet structure, in which the β-sheets are perpendicular to, and intermolecular hydrogen bonds are parallel to the self-assembled fibers. ^8, 9^ The unique characteristic of this conformation to accommodate tightly packed interactions is reflected in the robust enzymatic and mechanical stability of amyloids which has inspired the generation of amyloid nanomaterials.^10-12^ Our group has previously demonstrated that hydrophobins, a class of functional amyloids, present different morphologies and mechanical properties depending on temperature, solvent, pH, and salinity.^13^ The self-assembly process of amyloid proteins is known to be highly dependent on the protein or peptide sequence as well as its environment. ^14-18^ Because of these robust properties, there has been increasing effort toward developing design guidelines for peptide self-assembly in order to exploit their functional properties for biomaterial applications. Godin and coworkers reported that modification of the Asn-21 residue in the islet amyloid polypeptide (IAPP) with amino acids varying in hydrophobicity (Gly, Pro, or Aib) significantly affected the intra- and inter-molecular interactions during self-assembly. Their results indicate that site-specific modifications either promoted amyloid fibril formation or modulated fibril formation pathway towards oligomeric species. ^19^ Furthermore, amyloid nanowires with enhanced conductivities have been generated using engineered CsgA peptides containing polytyrosines or tyrosine-tryptophan mediated by π-π stacking.^20^ Nanowire formation of different thicknesses and packing densities has also been shown using CsgA, CsgB, or CsgA-CsgB fibers self-assembled on gold surfaces by Onur et al.^21^ They reported that nanofiber network morphologies strongly influenced biofilm optical properties, however, the structural features associated with fibril morphology were not investigated. In addition to the investigation of functional properties of naturally occurring amyloids, amyloid fibers have been generated from designed peptides to be applicable in several areas such as carbon dioxide adsorption, retroviral gene transfer, and artificial melanosome function.^22-24^ Although there has been some progress in identifying the functional properties of amyloid fibers, there is a dearth of understanding of the sequence grammar on the assembly process and structure–material property relationships necessary for designing *de novo* amyloid peptides for well-defined nanostructured biomaterials. Amyloidogenic core sequences that are only a few amino acid residues in length are strongly associated with the formation of the cross-β sheet structure;^25-27^ therefore, the investigation of short peptide sequences is a promising approach to serve as a model for broader applications.

Herein, we specifically examine differences in the self-assembled structures formed from *de novo* synthetic decapeptides (DPs) containing an amyloidogenic core and systematically varied single amino acid residues, designed to demonstrate individually the effects of polarity, hydrophobicity, and charge. The assembly process, secondary structure, fibril morphology, and nanomechanical properties are evaluated using atomic force microscopy (AFM), circular dichroism (CD), Fourier transform infrared spectroscopy (FTIR), and quantitative nanomechanical mapping (QNM). Relationships are established between de novo peptide amino acid sequence and secondary structure, fibril formation propensity, and nanomechanical properties for the creation of design rules for the development of tunable functional biomaterials.

## 2 Experimental methods

### 2.1 Materials

Ethyl cyanoglyoxylate-2-oxime (Oxyma), CL-MPA ProTide Resin (LL), and Fmoc protected amino acids were purchased from CEM peptides (USA). Trifluoroacetic acid (TFA), N-dimethylformamide (DMF), dichloromethane (DCM), diethyl ether, N.N’-diisopropylcarbodiimide (DIC), triisopropylsilane (TIS), ethane-1,2-dithiol (EDT), acetonitrile (MeCN) and all other solvents and reagents were purchased from ThermoFisher Scientific (USA) or Sigma-Aldrich Corporation (USA) at highest purity unless otherwise stated.

### 2.2 Peptide synthesis and purification

Decapeptides were synthesized using 9-fluorenyl methoxycarbonyl (Fmoc)-based solid phase synthesis on a Liberty Blue 2.0 automated microwave peptide synthesizer (CEM). Peptide synthesis was conducted on a 0.25 mmol scale using CL-MPA ProTide Resin (LL) (0.17mmol/g). Fmoc deprotection was achieved using (20 v/v%) piperidine/DMF. Coupling was carried out using Fmoc-protected amino acid (0.2 M), coupling reagent Oxyma (1 M), and DIC (0.5 M). After final amino acid coupling, the resin was washed with DCM followed by cleavage from the resin by stirring with a TFA/TIS/H_2_O/EDT (9.5:2.5:2.5:2.5) cleavage cocktail for 3 hours. Peptides were then precipitated using cold diethyl ether. The precipitate was centrifuged, the excess solvent was decanted, and the remaining dried pellet was then dissolved in 1:1 H2O:MeCN and lyophilized. Peptides were then purified using a Prodigy preparative reverse-phase HPLC (CEM) with a water/acetonitrile (each containing 0.1% v/v trifluoroacetic acid) gradient. The mass and identity of the eluting fractions containing the desired decapeptide were confirmed using electrospray ionization (ESI)-mass spectrometry (MS) on a ThermoFisher LXQ ESI-Ion trap mass spectrometer (Waltham, MA, USA). Sequences from the core-amyloid region of amyloid-β (Aβ) peptide were used as a positive control (DP(+)) while peptides made of random scrambled sequences rich in charged residues and negligible hydrophobic residues were used as a negative control (DP(-)). ^28, 29^

### 2.3 Fourier transform infrared spectroscopy (FTIR)

DP samples were prepared at ∼1.3 mM concentration by dissolving the dry peptides in 3:1 C_2_D_6_O:D_2_O, and FTIR data was obtained at room temperature. Samples were scanned for 1024 runs at 8 cm^-1^ resolution. Data were processed using OriginLab 8.0 after blank subtraction of 3:1 C_2_D_6_O:D_2_O spectra from the sample scan. Peak deconvolution was also done using OriginLab 8.0 using the Gaussian algorithm.

### 2.4 Circular dichroism (CD)

Far UV CD spectra (260-190 nm) for each decapeptide (DP) at a concentration of 30 µM in 3:1 ethanol/H_2_O was obtained by heating the sample from 10 to 90 °C in a 1 mm pathlength cuvette on a J-815 (Jasco, Easton, MD) spectropolarimeter. Scans were taken starting at 10°C with an average of 2 spectral scans at an interval of every 5 °C with a temperature gradient of 10 °C/ min and a delay time of 60 sec. Spectral smoothing using the Savitzky-Golay algorithm was applied to averaged spectra. In addition, far UV spectra of the samples after cooling from 90 °C back to 10 °C were obtained. Prior to each experiment, cuvettes were cleaned with 1% Hellmanex and blank 3:1 ethanol/H_2_O scans were done to confirm the cleanliness of the cuvette. The obtained data were processed using OriginLab 8.0 to calculate the mean residue ellipticity (MRE) and prepare the graphical plots.

### 2.5 Atomic force microscopy (AFM) and quantitative nanomechanical mapping (QNM)

Decapeptides were dissolved at 30 µM concentration in 3:1 EtOH: H_2_O and then sonicated at 75% power in a Misonix XL-2000 series sonicator for every 5 s with resting intervals of 30 s on ice. This cycle was repeated 20 times for each peptide sample. The presence of sonicated DP aggregates was confirmed by DLS using a Zetasizer DLS instrument by accumulating 15 scans per sample. Sonicated samples were immediately deposited on mica for further AFM analysis. All AFM experiments were performed using a Dimension Icon atomic force microscope (Bruker). AFM scanning was conducted using NanoScope 8.15r3sr9 software and the images were analyzed in NanoScope Analysis 1.50 software. Imaging was performed using a sharp silicon nitride cantilever (RTESP-300, nominal tip radius 8 nm; nominal resonance frequency of 300 kHz; nominal spring constant of 40 N/m) and a standard probe holder under ambient conditions with 512 × 512 data point resolution. AFM height and phase images were obtained simultaneously using tapping mode following a previously published procedure.^30^ A mica surface was cleaved using tape and then attached to a magnet. The mica was then treated with 150 μL of 3-aminopropyltriethoxysilane (APTES) solution (500 μL of 3-aminopropyltriethoxysilane in 50 mL of 1 mM acetic acid) for 30 min. The APTES solution was then decanted, and the mica substrate was rinsed three times with DI H_2_O, dried with N_2,_ and stored for an hour before use. The decapeptide solution (150 μL of 50 μM in 3:1 ethanol/water) was deposited on the mica surface and allowed to absorb for 30 minutes. The sample solution was decanted from the mica surface, and the mica was washed with 150 µL of di H_2_O three times, dried with N_2,_ and stored in a desiccator until imaging. Peak Force Quantitative nanomechanical mapping mode (PF QNM) was used to analyze the modulus of the peptide fibrils. Prior to each experiment, calibration of the cantilever deflection sensitivity was performed on a hard sapphire surface and the spring constant was determined using the thermal method. The tip radius was determined from the analysis of a roughened titanium surface (RS-15). The Young’s modulus of the peptide fibrils was obtained by analyzing nanomechanical maps that are produced by fitting the Derjaguin–Muller–Toporov (DMT) model to each force-indentation curve generated at each pixel scanned.^31^

## 3 Results and discussion

### 3.1 Rational design and characterization of *de novo* decapeptides

To generate customizable amyloid biomaterials, it is imperative to develop first principles on sequence grammar and amino acid priorities for amyloid formation. We examined the statistical prevalence of specific amino acid types (aliphatic, aromatic, hydrophilic, or charged) within the six amino acid stretches of amyloid-forming proteins in the database, and generated a library of *de novo* decapeptides (DPs, 10 amino acids in length), keeping the outer residues consistent and only making alterations to the core six residues in the sequence. Specifically, we kept the aliphatic residue, I at the 5^th^ position constant and varied the 6^th^ position with aromatic (F; DPII) and positively charged residue (K; DPIV). Similarly, we kept V at the 6^th^ position constant and varied the 5th position with polar residues (T; DPII and S; DPIII). The amyloid-forming propensities of the designed peptides were confirmed using the computational tool ZipperDB, which predicts the fibril-forming six amino acid stretches.^32^ Typically a Rosetta free energy score of less than -24 kcal/mol is reflective of amyloid-forming propensity as in the case of positive control, DP(+) (Table 1). ^32, 33^ All our designed peptides showed free energy less than this value but for the negative control (Table 1).

**Table 1.**
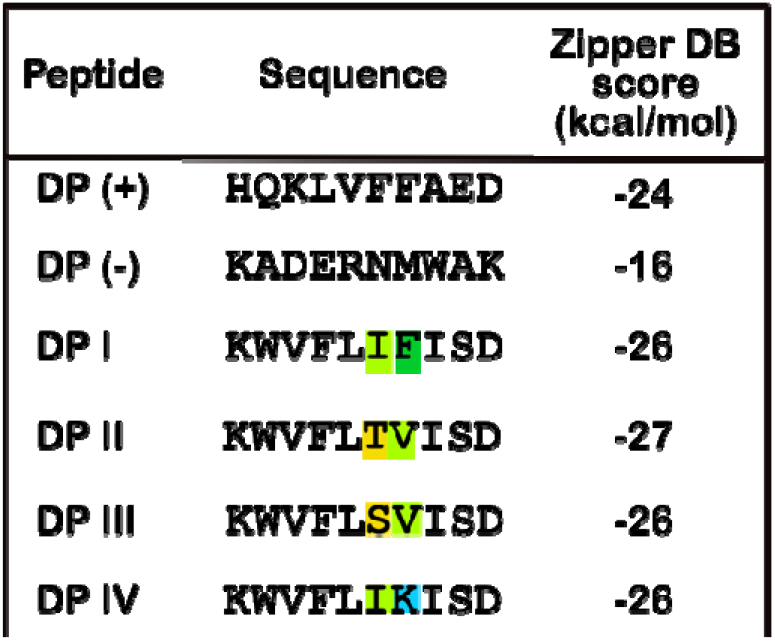
Decapeptide sequences with corresponding rosetta energies determined from ZipperDB.

The four decapeptides were synthesized via solid phase peptide synthesis (SPPS) (Fig S1), as described in the experimental section. Structures of positive and negative controls are given in Fig S2. Electrospray ionization mass spectrometry (ESI-MS) was used to confirm molecular weights and successful synthesis and deprotection for each peptide (Fig S3). The ESI-MS spectra and resulting m/z values were consistent with the expected mass distributions for each decapeptide sample.

### 3.2 Foldability of the peptides as a measure of their stability

We first sought to probe the stability of these peptides by subjecting them to denaturants such as SDS and guanidinium hydrochloride but could not see a response perhaps due to their highly aggregated state (data not shown). Therefore, we used the helix-inducing chaotrope, trifluoroethanol (TFE) to observe the conformational transition to α-helix as a function of temperature.^34^ Therefore, peptides at 30 µM concentration were dissolved in 100% (v/v) TFE and the conformational changes were monitored on a far-UV circular dichroism (CD) spectrometer by increasing the temperature from 10 to 90 °C at a rate of 10 °C/min. DPII, DPIII, and DPIV showed a random coil structure, indicated by the minima at 200 nm, immediately upon dissolving into TFE at room temperature (Fig 2 b-d). Surprisingly, TFE failed to induce α-helix, and the random coil structure persisted even with increasing temperature (Fig 2 f-h).

**Fig 1.**
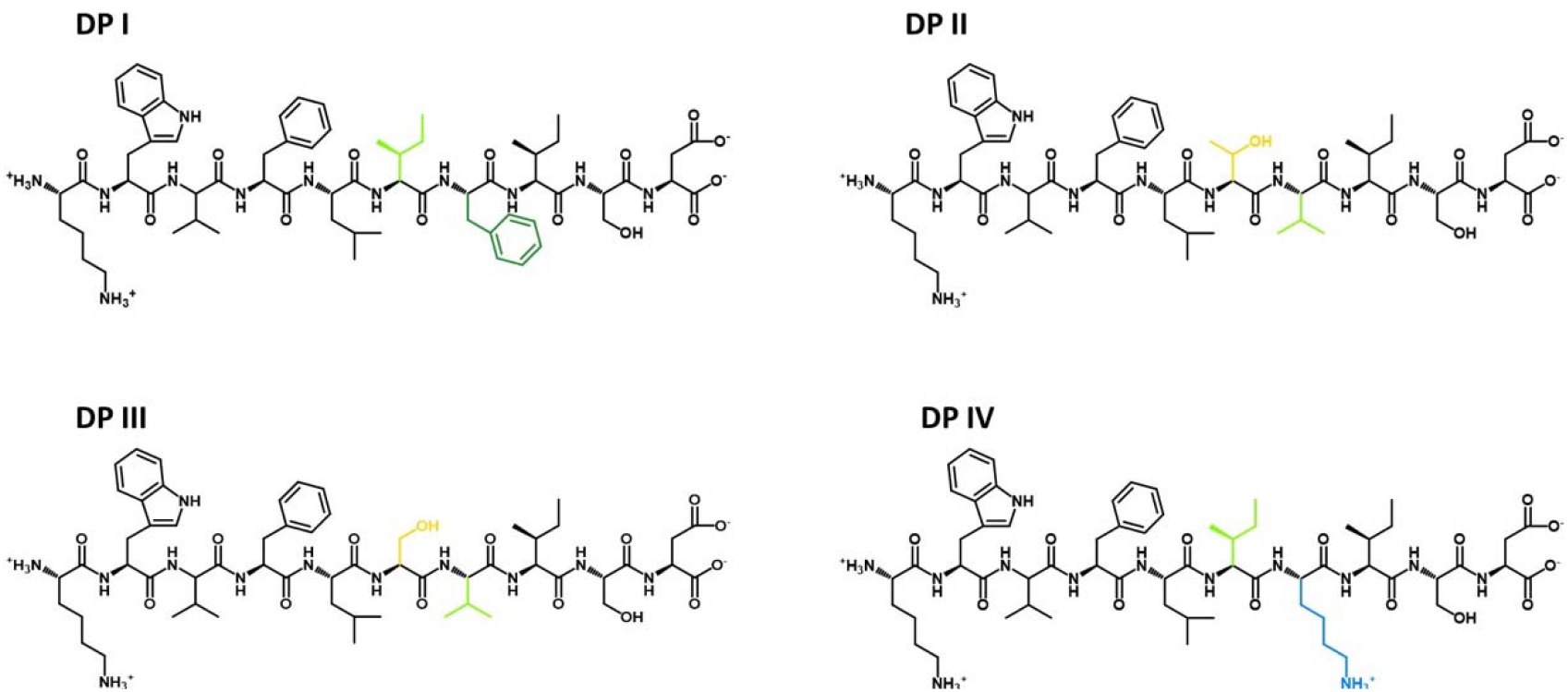
Chemical structures of the four designed decapeptides.

**Fig 2.**
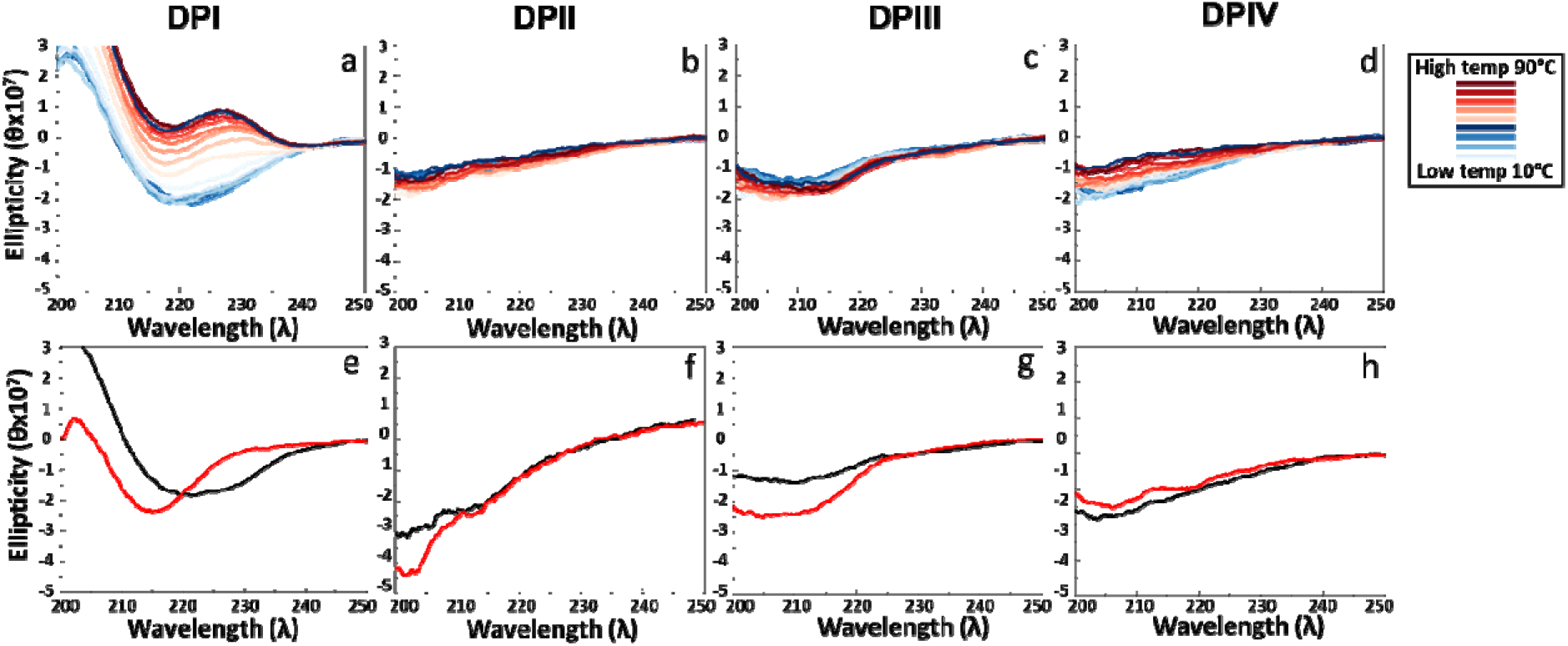
(a-d) Far-UV CD spectra as a function of temperature (from 10 – 90°C) for 30 µM DPI, DPII, DPIII, and DPIV, respectively, in TFE. (e-h) Far-UV CD scans at room temperature immediately after resuspension in TFE (black; pre-melt) and after the end at 90°C and equilibrating back at room temperature (red; post-melt).

DPIII, however, showed a transition to a β-sheet to a small extent, indicative of a possible ensemble with mixed conformers (Fig 2c and g). In contrast, DPI showed distinct conformational transitions with increasing temperature (Fig 2a). Unlike the other DPs, upon addition of TFE, DPI showed a transition to a spectrum with a minimum at 220 nm, indicative of a β-sheet but not to an expected α-helix, suggesting that the peptide could be resistant to the transition owing to its stability (black; Fig 2e). Surprisingly, even upon increasing temperature, the conformation did not change to a helix but toward type IV β-turn, alternatively termed as distorted or hybrid variants of turns I (III) and I’ (III’) β-turn with a positive maximum at 222 nm wavelength.^35^ The post-melt spectrum at room temperature showed a minimum around 215 nm indicative of an ensemble with random coil and β-sheet conformations. It has been shown previously that interactions between aromatic and aliphatic residues can drastically affect the stability of peptide conformations in TFE, and an interplay between hydrophobic and aromatic π-π interactions can act as a driving force for self-assembly.^36^ Therefore, it is likely that the substitution of Val residue to a Phe in the sixth position and having an aliphatic hydrophobic residue, Ile, in the fifth position in DPI plays a major role in the increased resistance for conformational conversion to an α-helix in TFE. It is also equally intriguing that peptides DPII, III, and IV failed to show any conformational transition and failed to adopt any secondary structure. This could mean that either these peptides form aggregates by a network of non-specifically aligned molecules, or the introduction of polar residues in the fifth or sixth positions in the peptide (DPII, DPIII, and DPIV) greatly destabilizes the ability to self-associate.

### 3.3 Secondary structures of the peptides in ethanol

To obtain insights into the secondary structures of the peptides solvated in ethanol, FTIR analysis was performed. DPs were dissolved in 3:1 deuterated ethanol (C_2_D_6_O): D_2_O at a concentration of 1.3-1.5 mM and were subjected to data collection (see details in the experimental section). DPI showed a deconvoluted spectrum containing three distinct peaks at 1625 cm^-1^, 1677 cm^-1^, and 1685 cm^-1^. While 1625 and 1685 cm^-1^ peaks correspond to an antiparallel β-sheet, the 1677cm^-1^ peak corresponds to a turn (Fig.3a).^37, 38^ The peptide DPII showed two peaks at 1618 cm^-1^ and 1675 cm^-1^, which are also indicative of parallel β-sheet and turn conformations, respectively (Fig. 3b). Similar aggregated parallel β-sheet and turns were also observed for DPIII, indicated by peaks at 1620 cm^-1^ and 1677 cm^-1^ (Fig. 3c).^39^ The FTIR spectrum for DPIV indicates a mixed conformation with antiparallel β-sheets with peaks at 1620 and 1690 cm^-1^, a peak representing disordered structure at 1640 cm^-1^, and a small percentage of a peak representing turn conformation at 1677 cm^-1^ (Fig. 3d).^40^

**Fig 3.**
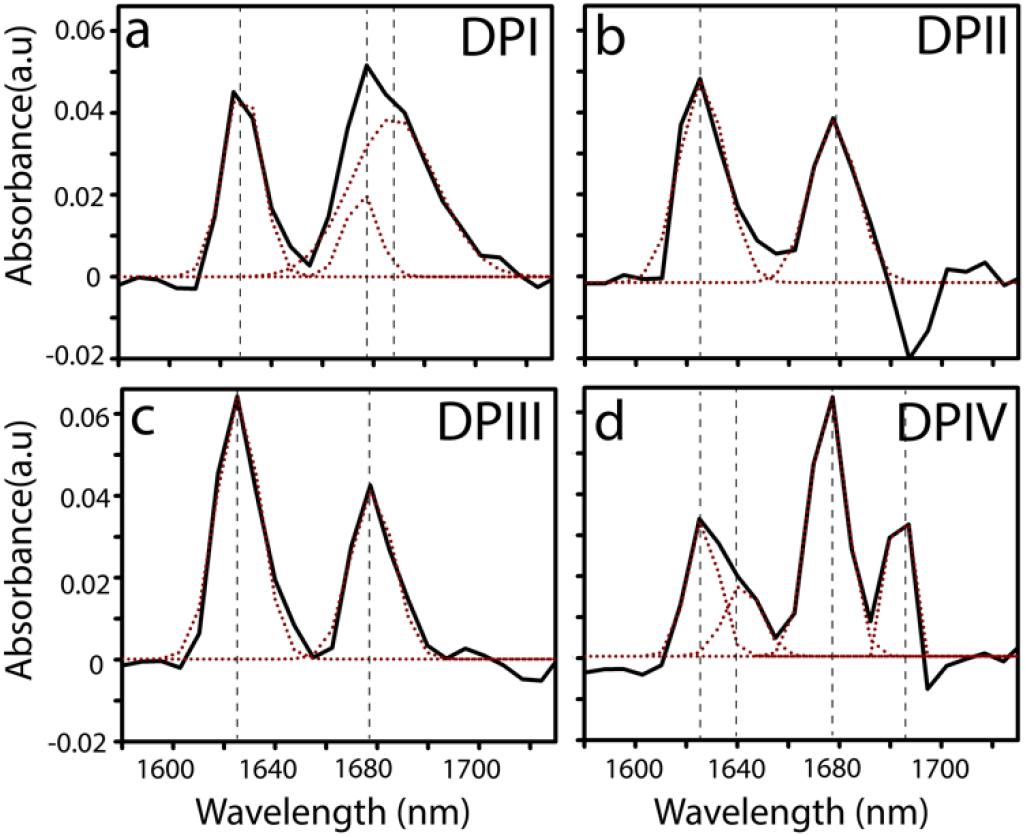
(a-d) FTIR spectra of DPI, DPII, DPIII, and DPIV peptides in 3:1 ethanol-water solvent. Black lines indicate the raw spectra and the dotted lines indicate the deconvoluted spectra.

### 3.4 Effect of temperature on peptide conformation and morphology

In order to probe changes in peptide structure as a function of temperature without the influence of any chaotropic agents, 30 µM peptide samples dissolved in 3:1 EtOH:H_2_O were analyzed by far UV CD spectra by increasing the temperature at a rate of 10 °C/min from 10 to 90 °C similar to the TFE experiment.

DPI, which is the most hydrophobic of all the peptides investigated, at lower temperatures showed a minimum at 215 nm and a shoulder at 226 nm suggestive of a helical structure (Fig 4a).^41^ These minima are red-shifted from canonical α-helix, which could be attributed to the coupling of carbonyl transition dipoles in a closely packed arrangement in the aggregated peptide. However, as the temperature increases over 50 °C, shifts towards a negative minimum at 222 nm and a positive maximum approaching 200 nm were observed, indicating a change in conformation towards a type II β-loop, as previously observed (Fig 4a).^42-44^ A similar trend was observed in the case of DPII with a transition from 215 and 222 nm to 225 nm (Fig 4b), indicating a transition from α-helix to type II β -loop conformation with an increase in temperature. Interestingly, a similar but very subtle transition from a bimodal distribution at 225 and 215 nm to a unimodal minimum towards 222 nm was observed upon increasing temperature in DPIII (Fig 4c), most likely indicating a mixture of α-helix and β-sheet conformations within the peptide sample. The similarity in the behavior of the peptides DPII and DPIII can be appreciated by the fact that the two only differ subtly by one amino acid residue at position six, where DPII has a slightly more polar Ser residue compared to the Thr residue in DPIII. It has been seen previously that interchanging serine and threonine usually has little effect on the resulting peptide secondary structures (α helix or β sheet), however, the differences in hydrogen bonding and hydrophobicity can affect solvent interactions resulting in differences in side chain residue orientations.^45, 46^ In the case of DPIV, only a negative minimum at 200 nm was observed indicating a random coil conformation^47^, and no change in this conformation was observed with increasing temperature, similar to the peptide’s behavior in the presence of TFE. The CD data for DPI, DPII, and DPIII agree with the FTIR data confirming the heterogenous mixture of structures (Fig 4a-c). However, while the CD spectrum for DPIV in ethanol-water mixture showed predominantly random coil conformation (Fig 4d), FTIR showed an ensemble containing a combination of secondary structures (Fig 3d). CD and FTIR are complementary techniques; while CD provides a time-averaged conformation of ensembles, FTIR could provide qualitative estimates of secondary structural elements present. Therefore, one could say that the combination of secondary structures detected by FTIR is accurately captured by the time-averaged bulk random coil conformation by CD.

**Fig 4.**
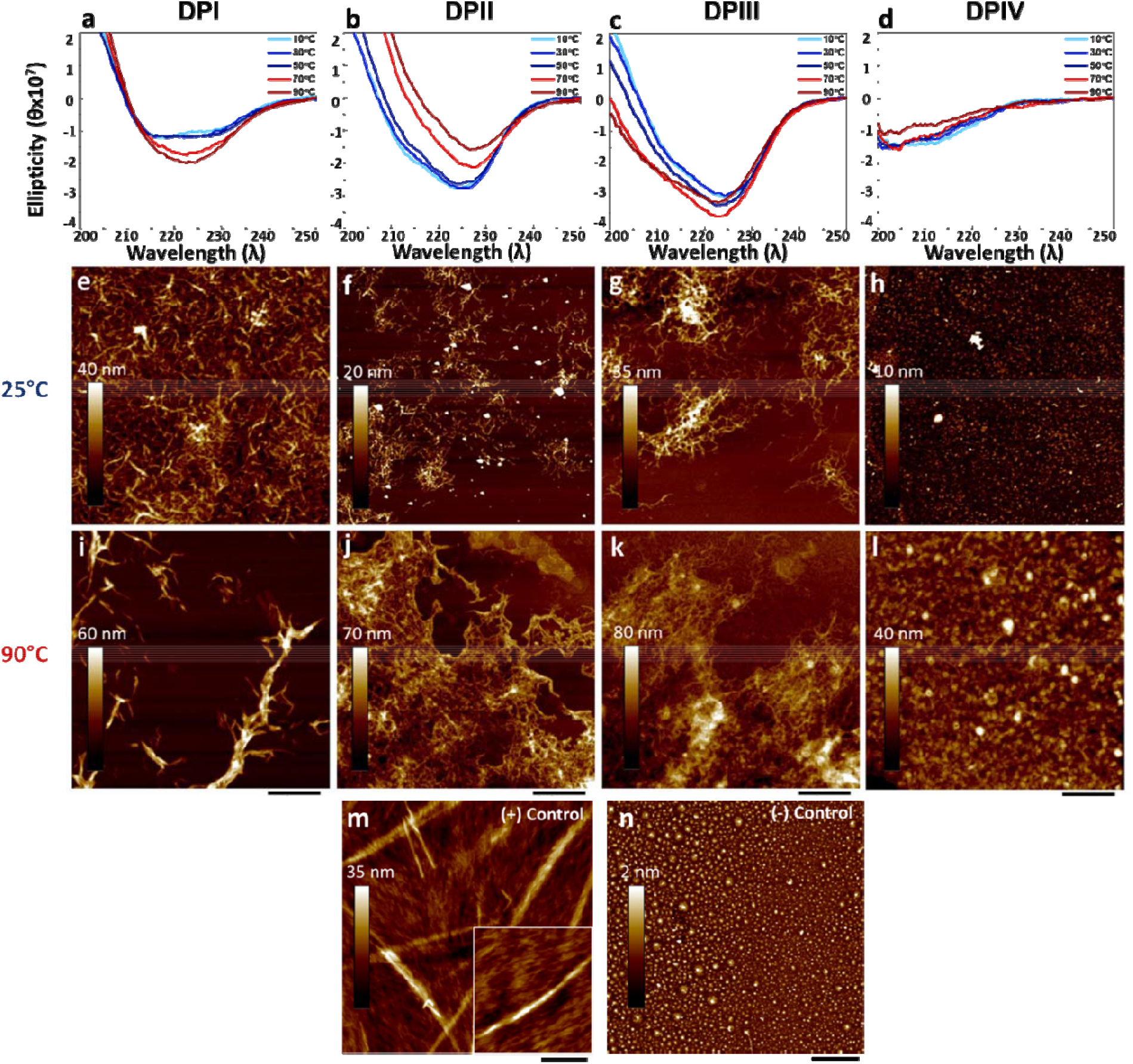
(a-d) Far-UV CD spectra at the indicated temperatures for 30μM DPI, DPII, DPIII, DPIV, respectively, in 3:1 ethanol:water. (e-l) AFM tapping mode height images of the samples from (a-d) following incubation at 25 °C (e-h) and 90 °C (i-l). (m-n) The positive control, DP(+) shows the expected fibril like morphology whereas the negative control, DP(-), shows the absence of fibril formation. All AFM images are 5μm x 5μm except for the inset in (m) which is 1μm x 1μm.

AFM tapping mode was used to image the peptides to further probe morphological changes as a function of temperature. Aliquots of the peptide samples analyzed for CD were taken and processed for AFM analysis as detailed in the experimental section along with the controls, DP (+) and DP (-). As expected, the positive control, DP (+) showed fibrils while the negative control, DP (-)showed no organized features (Fig 4 m and n). The peptides, DPI, DPII, and DPIII showed fibril-like morphologies at both high and low temperatures, albeit with distinct differences (Fig 4e-g and i-k). DP I showed numerous short fibril-like structures at 25 °C (∼5 nm diameter, 10 – 20 nm in length), as well as large isolated fibrils at 90 °C (10-100 nm diameter, µm lengths), consistent with the increased organization indicated by CD. A caveat to be borne in mind is that since ethanol evaporates at 90 °C, the association of fibril structures observed could be due to a combination of temperature-based effects and those due to solvent evaporation.

Nevertheless, the fibrillar structures observed are consistent at both low and high temperatures. The peptide DPII displayed smaller and more dispersed fibrils at 25 °C, and a dense network of larger fibrils (µm length, 10 – 100 nm diameter) at 90 °C (Fig 4f and j). Aggregation at 90 °C showed a three-fold increase in the height scale (20 – 70 nm), which could be due to association due to evaporation. DPIII behaved similarly to DPII, with small dispersed fibrils at 25 °C, and larger aggregated features, and increased height scale, from 35 to 80nm at 90 °C (Fig 4g and k). Higher resolution images of DPII and DPIII at 25 °C showed helical-like features (Fig 5a and b), consistent with the heterogeneous mixture of secondary structures observed with circular dichroism in Fig 2a and b. However, DPIV showed no defined structures at low and high temperatures, which is in agreement with the random coil structure indicated by CD (Fig 2d). It is likely that the addition of a Lys residue and the positive charge in DPIV increases the solubility of the peptide, thereby hindering and inhibiting aggregation.

**Fig 5.**
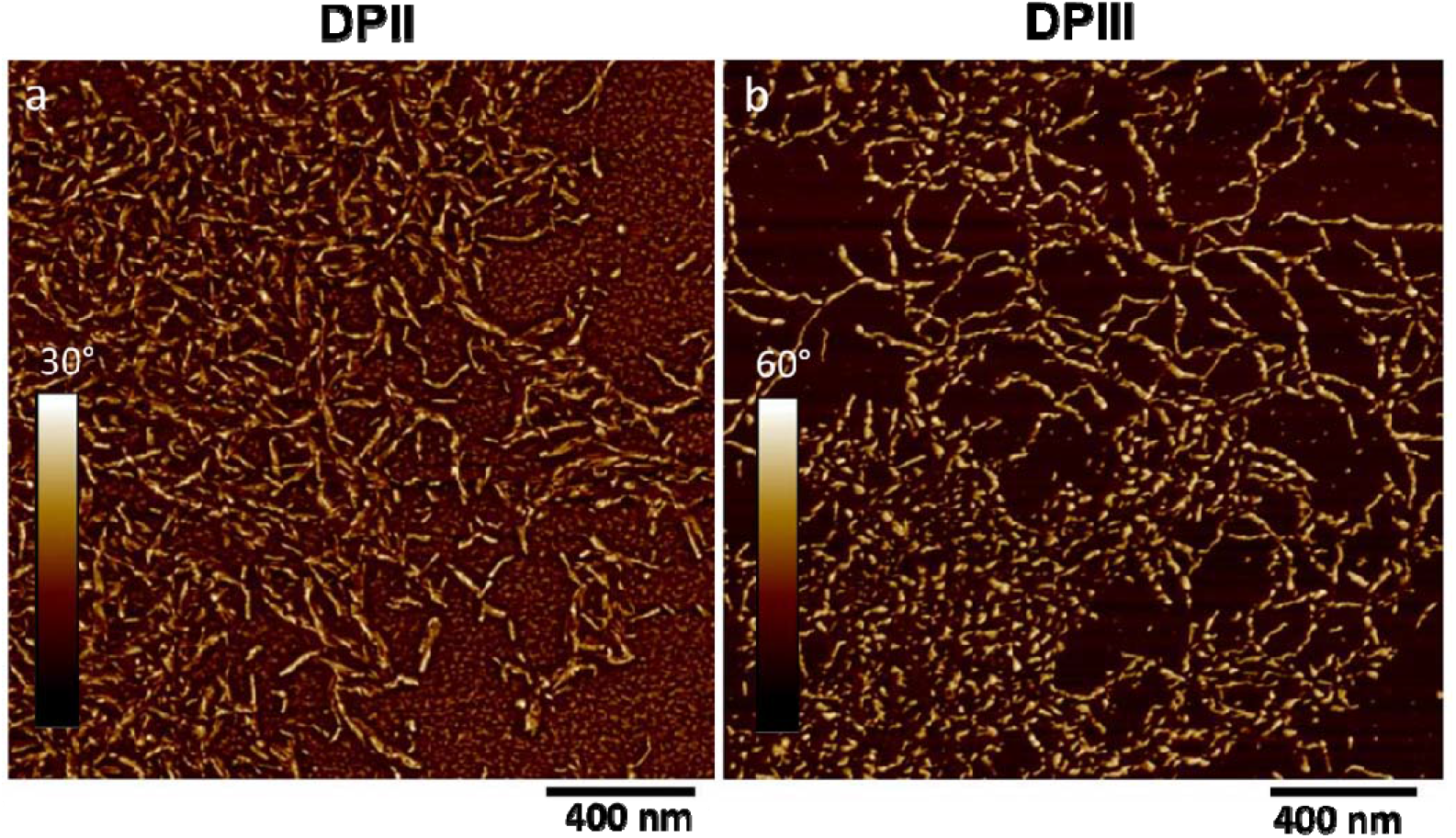
AFM Phase images of DPII (a) and DPIII(b) after incubation at 25°C. Both peptides exhibit subtle differences in a twisted fibril-like morphology.

### 3.5 Nanomechanical properties of the peptides show distinct differences

AFM PeakForce-QNM was used to investigate fibril nanomechanical properties of the peptides and correlate them with their sequence. PeakForce-QNM allows the measurement of height, adhesion, nanoindentation, and DMT modulus simultaneously.^31^ Nanomechanical properties determined from force measurements in AFM-QNM may be related to structural differences that are not evident from topographical information alone. It has been demonstrated that amyloid fibril mechanical properties are strongly correlated to their secondary structure content, where an increase in β-sheet structures increases rigidity, and an increase in α-helices results in reduced rigidity.^12, 48, 49^ For this study, the peptides were prepared by dissolving samples at 30 µM in 3:1 EtOH: H_2_O, sonicating to disperse fibrils, and immediately casting onto mica for analysis. Following imaging, force data were collected along the longitudinal axis of isolated fibrils, and modulus was obtained via the DMT method as described in the experimental section. Nanomechanical data for DPIV was not obtained, as it did not exhibit fibrillar morphology. Figs 6 – 8 show representative AFM height and modulus images of DPI, DPII, and DPIII. Two different fibril types with different modulus behaviors were observed for DPI. One fibril type appeared rough with irregular features and measured modulus values varying from 3 – 50 GPa (Fig. 6e, line 1 and 2), whereas the other appeared homogeneous with more consistent modulus values of 2 – 5 GPa (Fig 6e, line 3 and 4). These observations indicate the heterogeneous nature of the assembled peptides. The high modulus values may be attributed to DPI aggregates of higher order associated with the addition of a Phe residue, which has been shown to promote β-sheet formation and higher rigidity through π-π stacking and tight intermolecular packing.^50^ DPIII exhibited distinct helical-like morphologies with large modulus variation along the fibril longitudinal axis (Fig 8). The extreme peaks and valleys in the modulus values observed in individual fibrils (3 – 25 GPa) further confirmed the heterogeneity of the structures and indicated the presence of a mixture of turns and α-helices. DPII exhibited less defined fibril-like aggregates with lower and more consistent modulus values than those observed for the other two peptides (Fig 7). The lower rigidity seen in DPII compared to DPIII could be attributed to the extra methyl group on the Thr amino acid residue. It has been shown that methyl groups can act as plasticizers during protein self-assembly, increasing the overall flexibility of the peptide chains and affecting the thermal stability of the resulting secondary and tertiary structures.^51, 52^ All three decapeptide samples, however, demonstrated relatively stiff modulus values well within the range of previously reported amyloid-like fibrils. ^50, 53, 54^

**Fig 6.**
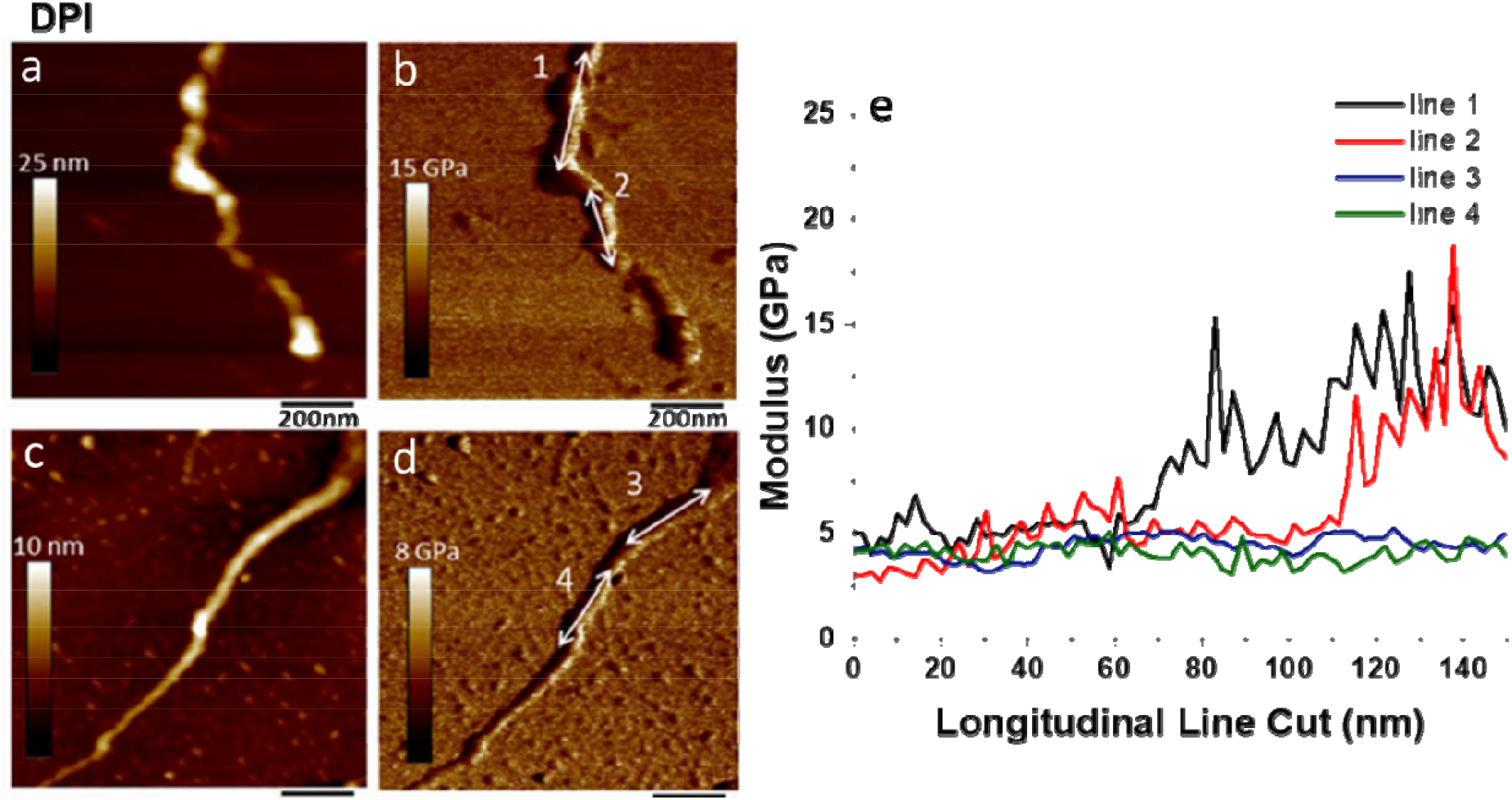
AFM Height (a, c) and DMT modulus (b,d) images of DPI. Modulus data of decapeptide fibrils obtained from longitudinal line cuts of DMT modulus data indicated by line number (e).

**Fig 7.**
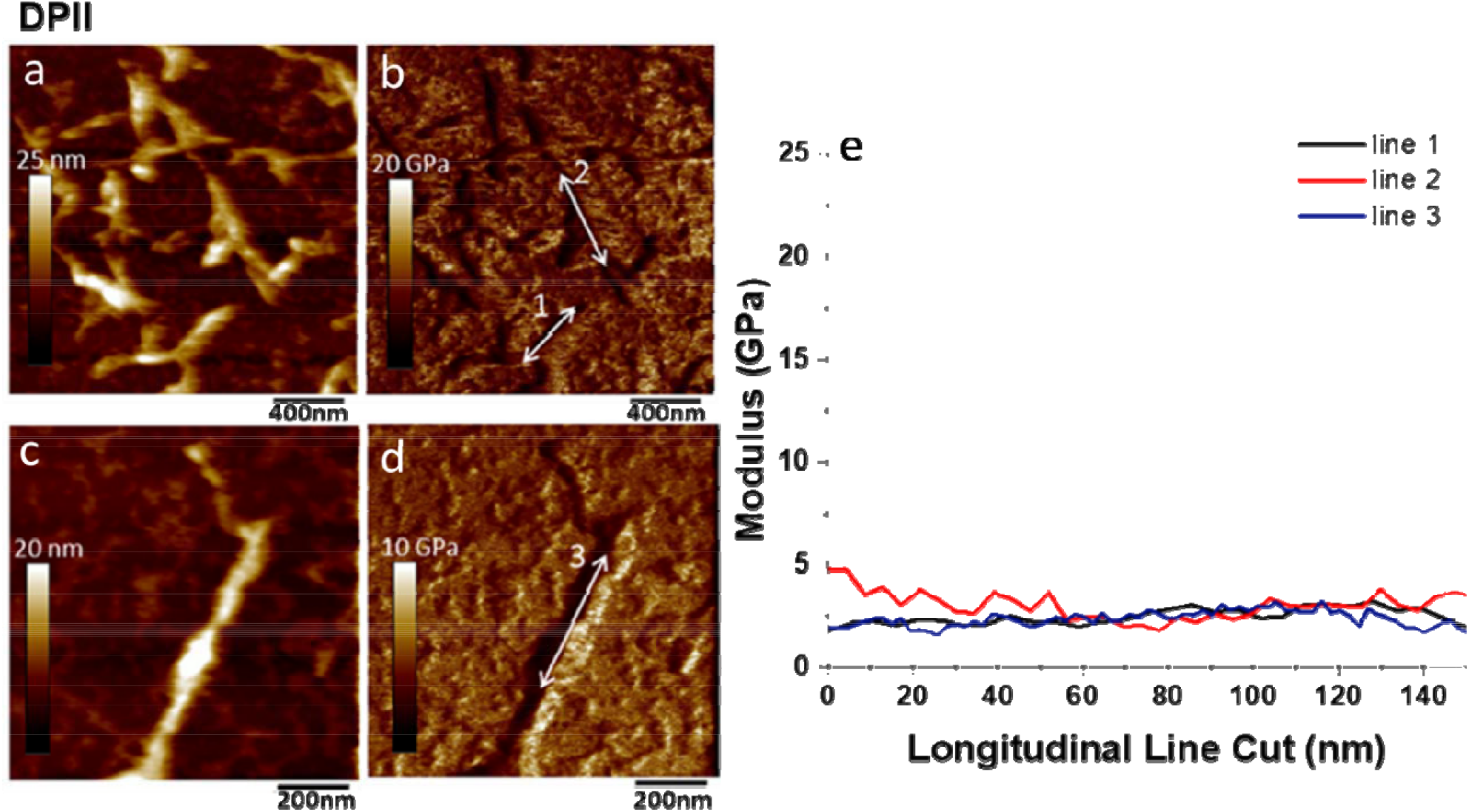
AFM Height (a, c) and DMT modulus (b,d) images of DPII. Modulus data of decapeptide fibrils obtained from longitudinal line cuts of DMT modulus data indicated by line number (e).

**Fig 8.**
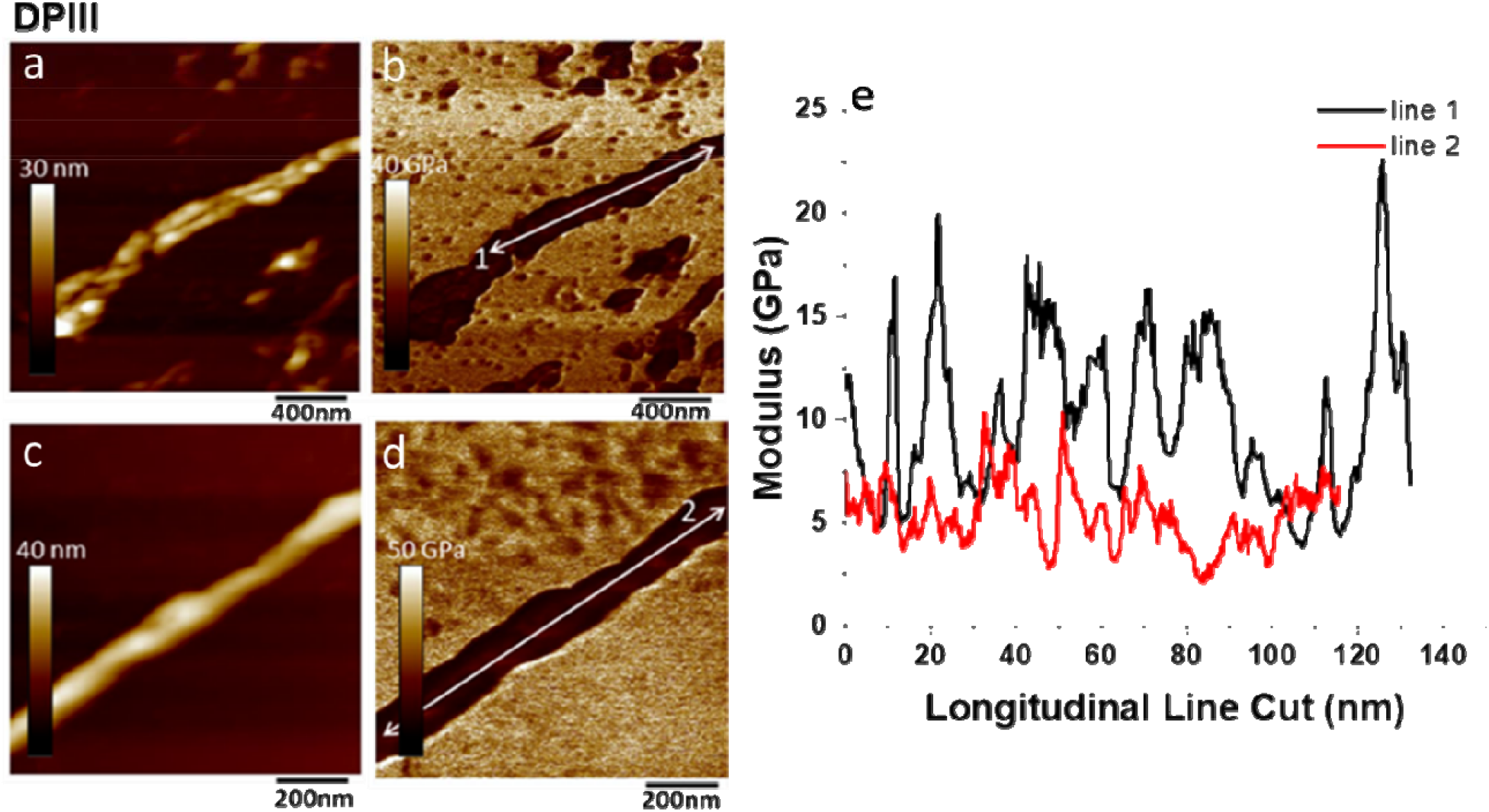
AFM Height (a, c) and DMT modulus (b,d) images of DPIII. Modulus data of decapeptide fibrils obtained from longitudinal line cuts of DMT modulus data indicated by line number (e).

## 4 Conclusions

The data presented here brings forth the sensitivity of peptide amyloid structures for subtle positional and chemical variations within the sequence, measured by their biophysical, morphological, and nanomechanical properties. More importantly, these studies provide cues for the efficient design of tailorable amyloid nanomaterials for a given application. The four peptides investigated here are part of a larger library of peptides that are specifically selected to investigate how subtle variations lead to the differences in their biophysical and mechanical properties. Specifically, we focused on positions 5 and 6 of the decapeptides and varied the chemistry from aliphatic hydrophobic, aromatic hydrophobic, polar, and charged (positive) amino acids. Although the lysine residue would carry a positive charge in neutral pH in buffers, in the ethanol-water solvent, it does not carry a charge but remains highly polar. As detailed above, we observed surprising differences in the aggregation propensity, structure, morphology, and nanomechanical properties of the fibrils generated. As expected, the most hydrophobic peptide, DPI showed robust amyloid properties; however, the peptide also showed the presence of turns alongside a β-sheet secondary structure which is intriguing considering the short length of the polypeptide. The other peptides too (DPII and DPIII), to varying degrees, showed the presence of turn conformations. At this point, it will be speculative to reason how the aggregation-prone short peptides contain turns and a high-resolution structural investigation by NMR in the future could provide answers. But parsimoniously one could note that the fibrils formed by the peptides are not pure cross-β sheet structures. The introduction of polar residues, Ser or Thr for one hydrophobic residue in the 5^th^ position (DPI and DPIII) results in observable changes in their morphologies and modulus. Substituting a much larger Lys residue in the 6^th^ position and retaining a hydrophobic residue in the 5^th^ in DPIV abrogates aggregation and fibril formation, suggesting that the 6^th^ position is intolerant to polar and possibly charged residues. In sum, we can conclude from this work that the core-amyloid regions in a polypeptide are not innocuous for subtle sequence variations as originally thought but are rather highly sensitive. It is imperative to gain an understanding of the positional preferences of amino acids within the amyloid core to be able to effectively design tunable amyloid-based nanomaterials.

## Author contributions

VR conceptualized the project with inputs from SEM, TDC, and PW. HGA and JS equally contributed to this work both intellectually and in experimental data collection. JS performed CD and FTIR analysis while HGA conducted AFM and modulus analysis. LKK helped with data analysis and manuscript writing. PW, TDC, and HGA performed peptide synthesis. VR, SEM, HGA, and JS contributed to manuscript writing while others were involved in editing.

## Acknowledgments

The authors would like to thank the National Science Foundation (2208349) for their financial support.

