## Supplementary material for "*De novo* amyloid peptides with subtle sequence variations differ in their self-assembly and nanomechanical properties": supp data

**Electronic Supplementary Information**

**Fig S1.** Scheme of decapeptide synthesis via solid phase peptide synthesis followed by cleavage from Wang resin, and deprotection of amino acids.

**Fig S2.** Structures of decapeptide positive (DP (+)) and negative (DP (-)) controls.

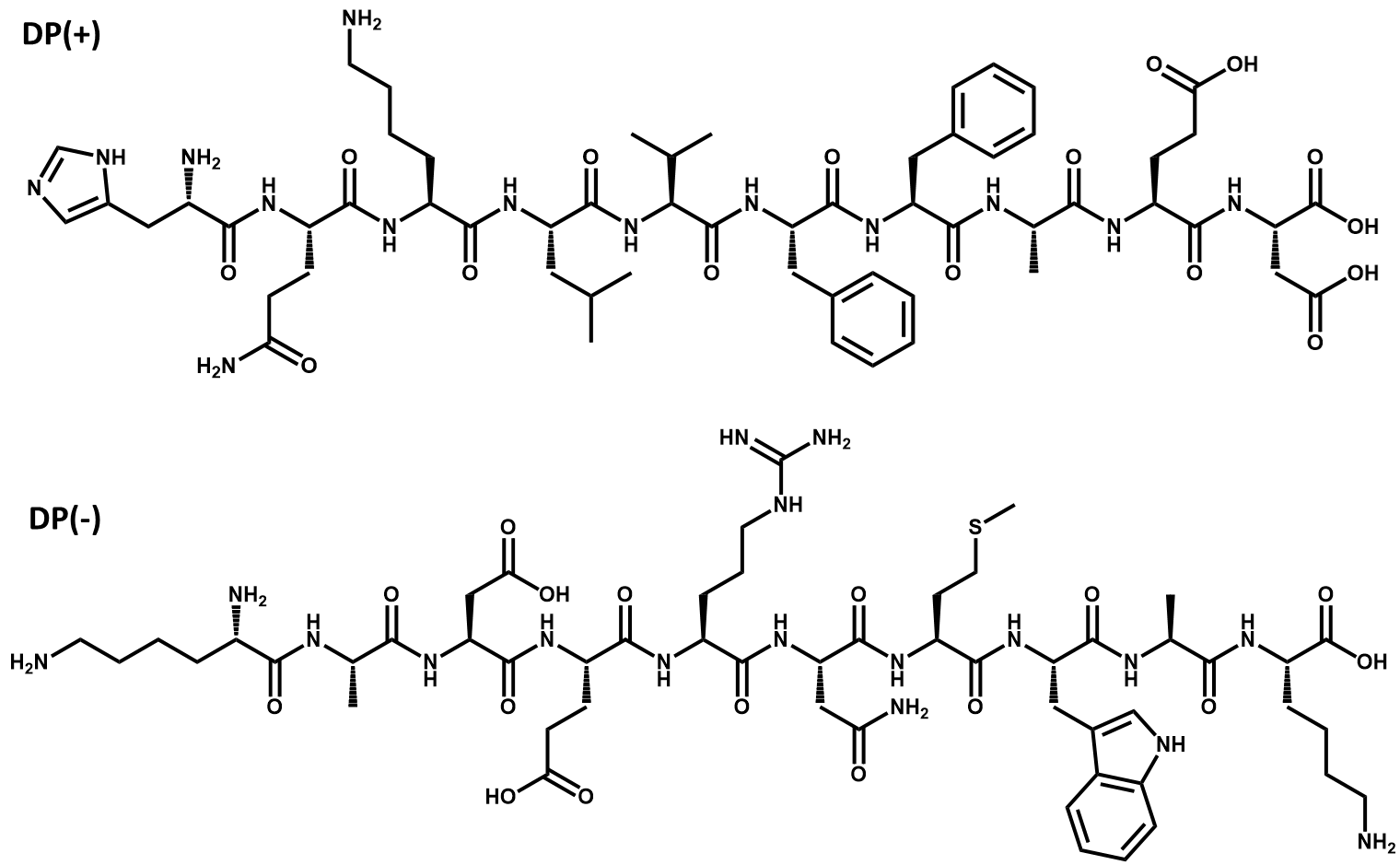

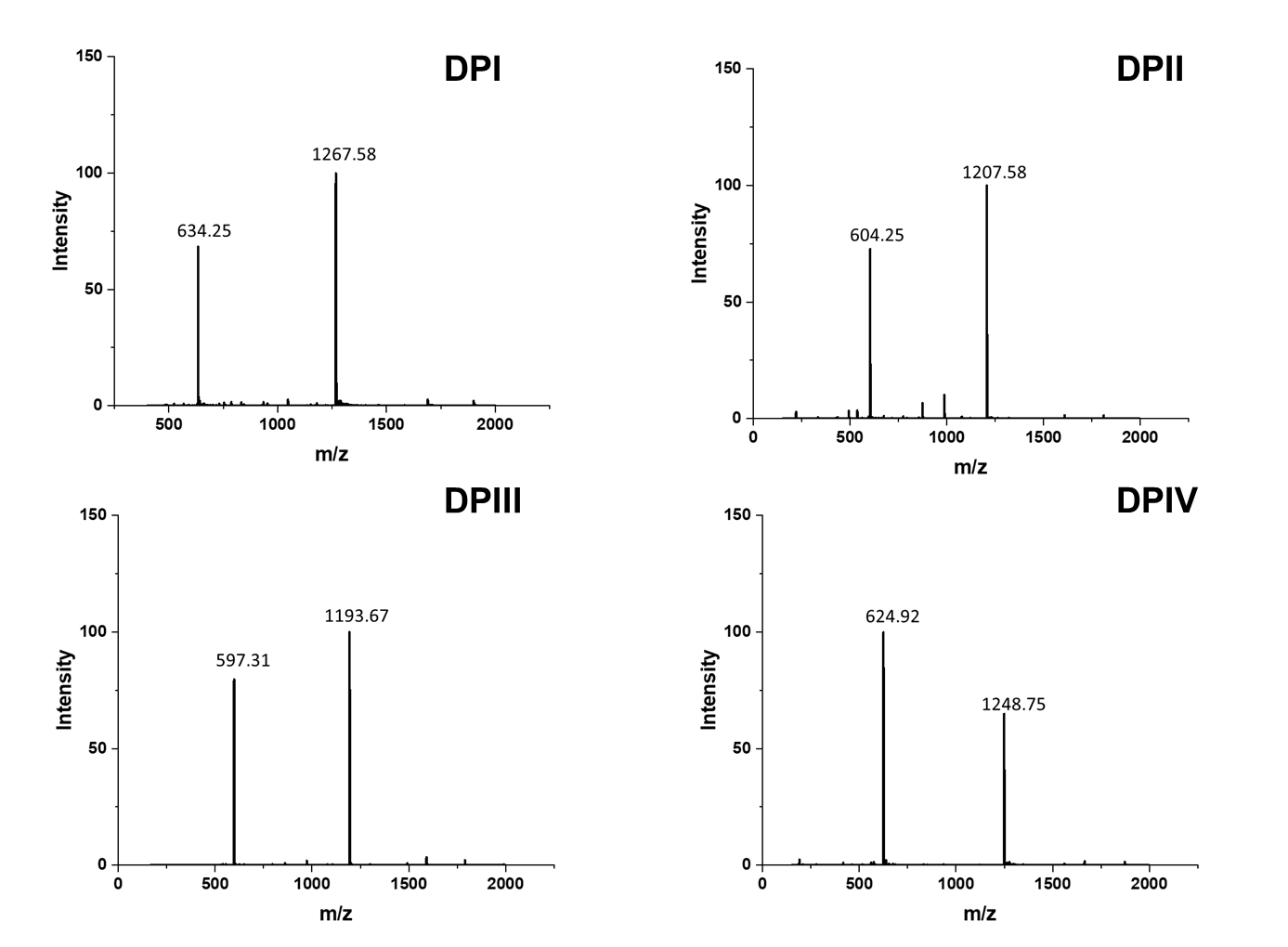
**Fig. S3** ESI Mass spectra of DPI, DPII, DPIII, and DPIV.
